# Mitochondrial PEX14 supports peroxisome biogenesis

**DOI:** 10.64898/2026.10.16.759530

**Authors:** Britta Nemeita, Anne Sophie Hammann, Suraj Karki, Tolga Soykan, Livia Wieland, Martin Lehmann, Sven Thoms

## Abstract

Peroxisomes are essential for cellular homeostasis and human health due to their central metabolic and physiological functions. The peroxisomal membrane protein PEX14 is indispensable for matrix protein import into peroxisomes. Recent studies support a hybrid model in which pre-peroxisomal vesicles originating from both, mitochondria and the endoplasmic reticulum, fuse to generate peroxisomes. PEX14 has repeatedly been observed at mitochondria in biogenesis disease models or upon protein overexpression, but such localization has generally been interpreted as aberrant mislocalization. Whether endogenous PEX14 naturally localizes to mitochondria and contributes to peroxisome biogenesis has remained unknown. Here, using high-content and super-resolution microscopy, we demonstrate that endogenous PEX14 localizes to the mitochondrial outer membrane. We show that ectopically expressed PEX14 accumulates at mitochondria, and PEX14 variants unable to access mitochondria and restricted to peroxisomes fail to rescue the PEX14 knockout phenotype. Vice versa, a mitochondria-targeted PEX14 variant supports peroxisome biogenesis. Our results identify the physiological mitochondrial pool of PEX14 in human cells and establish it as an essential intermediate in de novo peroxisome biogenesis.

## Introduction

Peroxisomes are ubiquitous eukaryotic cell organelles involved in lipid and reactive oxygen species (ROS) metabolism^1^. Recent work has involved peroxisomes in a broad array of physiological functions, including ferroptosis, immunity, inflammation, and ageing^2–6^. Peroxisomes have long been conceptualized as self-contained metabolic organelles, assembled through a largely autonomous biogenesis pathway^7^. In this model, the peroxisomal membrane proteins PEX13 and PEX14 form the core of the translocation machinery, onto which the receptor protein PEX5 docks to conduct the cargo into the peroxisome. PEX5 is subsequently ubiquitylated by a heterotrimeric E3 ligase complex and recycled by a peroxisomal AAA-type ATPase complex in conjunction with its membrane anchor PEX26^6,8–10^. The central ideas of the translocation mechanism are less controversial. However, the autonomy of the biogenesis proteins is increasingly difficult to reconcile with accumulating evidence that places peroxisomes within a highly interconnected organelle network^11,12^. While the distribution of metabolic pathways across multiple organelles involving peroxisomes is widely accepted^1^, the extent to which the endoplasmic reticulum (ER) and, more recently, mitochondria contribute to peroxisome biogenesis has remained debated for long.

Peroxisomes and mitochondria are closely linked at the metabolic level, functionally partitioning fatty acid oxidation. Peroxisomes mediate very-long-chain fatty acid (VLCFA) shortening and generate hydrogen peroxide, whereas mitochondria carry out complete oxidation coupled to ATP production. Beyond this division of labor, both organelles are integrated through reactive oxygen species (ROS) signaling, metabolite exchange at membrane contact sites^8,13–17^, and coordinated control of organelle dynamics and inheritance. Shared components involve the fission machinery, such as Drp1, Fis1, or Mff1^18–21^, other proteins with dual localization^22^, as well as the MAVS common signaling platform in antiviral responses^23^.

Under pathological conditions, particularly in peroxisome biogenesis disorders (PBDs) caused by inheritable defects in peroxins, peroxisomal membrane proteins, including biogenesis factors, can be redirected to mitochondria^24–27^. This mistargeting likely exacerbates cellular dysfunction beyond the primary peroxisomal defect^17,28–30^. Accordingly, mitochondria-associated peroxisomal proteins have typically been interpreted as aberrantly localized.

However, more recently, the identification of protein trafficking routes that transiently involve mitochondria has challenged this interpretation and prompted a re-evaluation of peroxisome biogenesis models. Work from the McBride group using PEX3-deficient fibroblasts demonstrated that peroxins such as PEX14 and exogenously expressed PEX3-YFP localize to mitochondrial membranes, from which functional peroxisomes can emerge^31^. Based on these findings, a hybrid model has been proposed in which peroxisome formation originates from both the ER and mitochondria^31^. According to this model, PEX16-positive pre-peroxisomal vesicles (ppVs) derived from the endoplasmic reticulum (ER) fuse with mitochondrial-derived vesicles (MDVs) containing PEX14 for de novo biogenesis of peroxisomes. MDVs themselves comprise a heterogeneous population of vesicular structures, typically smaller than 150 nm in diameter, that participate in multiple physiological processes, including mitochondrial quality control^32^. A recent study in neurons identified three distinct MDV populations differing in size, composition, and function, including a PEX14-positive population, supporting the model that peroxisomes can arise from both ER- and mitochondria-derived membranes^33^. Taken together, these findings suggest that PEX14 may also be involved during earlier stages of de novo peroxisome biogenesis. When incorporated into mature peroxisomes, PEX14 forms a complex with PEX13 at the peroxisomal membrane, which mediates the docking and translocation of the matrix protein import receptor PEX5^34,35^. The model of mitochondria-dependent peroxisome biogenesis pathways has recently gained support by the characterization of the E3-ubiquitin ligase MARCH5, which localizes to mitochondria and peroxisomes and assists MDV formation by directly interacting with PEX26^36–38^.

Direct evidence, however, that endogenous peroxins such as PEX14 naturally localize to mitochondria and that mitochondrially localized PEX14 actively drives peroxisome biogenesis is still lacking. Here, using confocal microscopy combined with high-content image analysis and dSTORM, we specifically investigated the localization and function of endogenous wild-type PEX14. We systematically analyzed the distribution of PEX14 within peroxisomal and mitochondrial membranes in human cells. Furthermore, by expressing distinct PEX14 variants, we demonstrate that preferentially mitochondria-targeted PEX14 supports de novo peroxisome biogenesis.

## Results

### Endogenous PEX14 localizes to the mitochondrial membrane

PEX14 has repeatedly been reported to localize to mitochondria^31,33^. However, these studies were often performed under pathological conditions caused by the knockdown or knockout of peroxins, which disrupts peroxisome biogenesis^25,31^. Not only depletion but also overexpression of peroxins, including but not limited to PEX14, can perturb their native subcellular localization and cause a spillover of peroxisomal membrane proteins into other cellular compartments^24,39^.

To investigate the physiological subcellular localization of PEX14 in relation to mitochondria, we used unmodified U2OS cells and employed confocal fluorescence microscopy (Fig. 1a). Using commercially available antibodies in combination with mitotracker staining, we examined the localization of peroxisomal proteins and mitochondria without perturbing either organelle through the exogenous expression of marker proteins. The use of primary antibodies raised in the same host species (rabbit) allowed the application of a single secondary antibody across all stainings, thereby ensuring maximal comparability between samples.

**Fig. 1.**
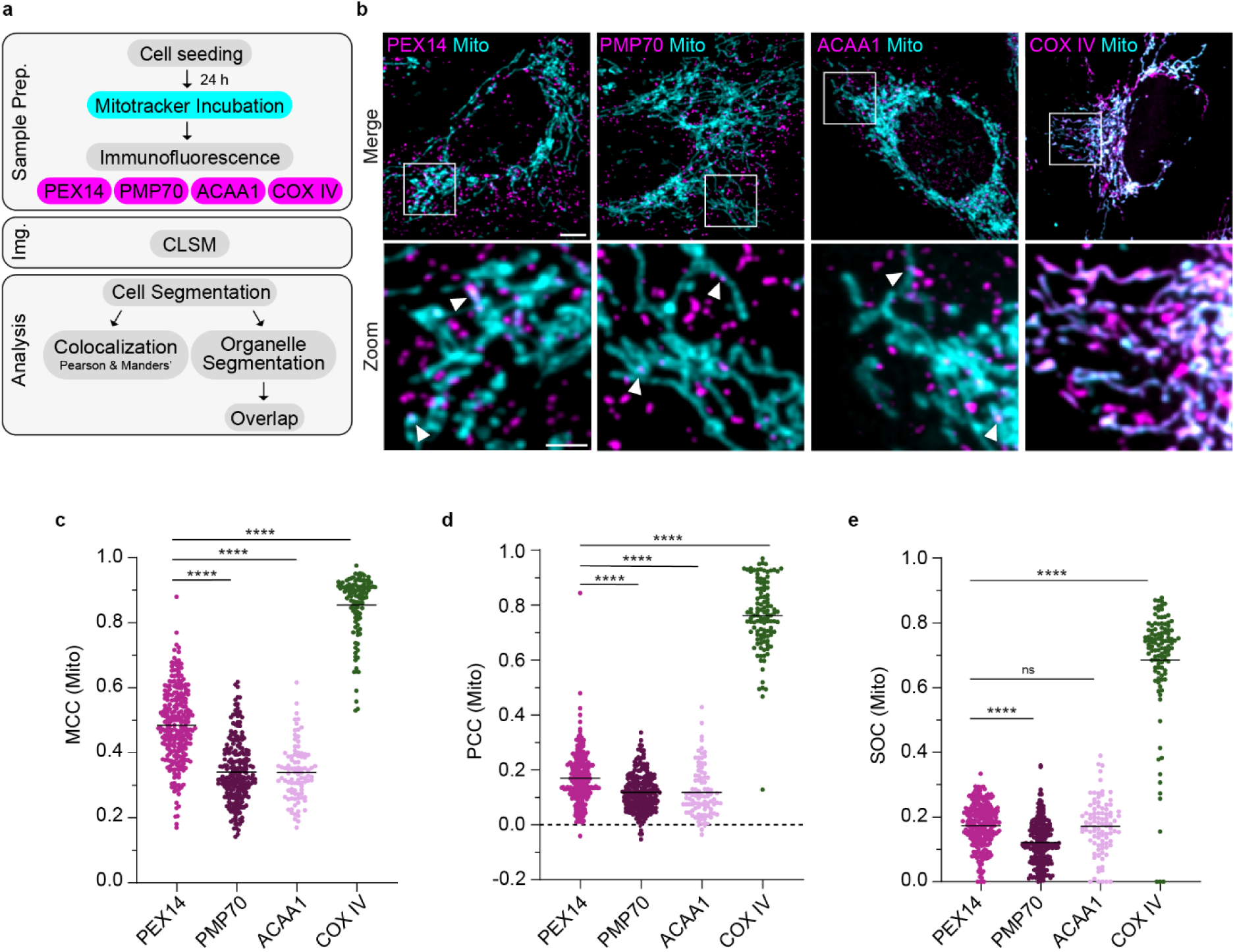
Endogenous PEX14 localizes at the mitochondrial membrane in U2OS cells. **a** Workflow of sample preparation for confocal imaging, including subsequent analysis of cells and organelles. **b** Representative two-color confocal fluorescence microscopy images of U2OS WT cells immunostained for endogenous PEX14, PMP70, ACAA1, and COX IV (magenta). Mitochondria were stained with mitotracker (cyan). Examples of colocalizing puncta are indicated by arrowheads. Scale bar: 5 µm. Zoom of the indicated box: scale bar: 2 µm. **(c-e)** Quantitative analysis of the colocalization of PEX14 (N = 261 cells), PMP70 (N = 223 cells), ACAA1 (N = 94 cells), and COX IV (N = 113 cells) with mitotracker. At least three independent experiments were conducted. Cells are represented as individual points. The central line represents the median. Statistical analysis was carried out using the Kruskal-Wallis test with two-sided Dunn’s multiple comparison test: ns > 0.05; *P ≤ 0.05; **P ≤ 0.01; ***P ≤ 0.001; ****P ≤ 0.0001. **c** MCC with mitotracker: PEX14: 0.48, PMP70: 0.32, ACAA1: 0.34, COX IV: 0.89. **d** PCC with mitotracker: PEX14: 0.17, PMP70: 0.11, ACAA1: 0.10, COX IV: 0.76. **e** SOC with mitotracker: PEX14: 0.18, PMP70: 0.12, ACAA1: 0.17, COX IV: 0.69.

Focusing on the distribution of PEX14, we observed distinct puncta throughout the cell. As expected for peroxisomal proteins, this pattern resembled that of the peroxisomal membrane protein PMP70 and that of 3-ketoacyl-CoA thiolase (ACAA1), a peroxisomal matrix protein localized within the peroxisomal lumen^40^ (Fig. 1b, Supp. Fig. 1). Several PEX14 puncta were located near mitochondria or colocalized with them (Fig. 1b, white arrowheads). These regions may represent membrane contact sites (MCS) between peroxisomes and mitochondria, ppVs, or proteins residing in the mitochondrial membrane.

To determine colocalization between peroxisomal markers and mitochondria, we quantified three complementary metrics reflecting distinct aspects of signal association. Manders’ colocalization coefficient (MCC) was calculated to determine the fraction of peroxisomal signal overlapping with mitochondrial signal, thereby providing a direct measure of signal co-occurrence. Pearson correlation coefficient (PCC) was used to evaluate the global correlation of fluorescence intensities, reflecting similarity in spatial intensity distributions independent of absolute signal levels. Additionally, we introduced a segmentation overlap coefficient (SOC), which calculates the relative area overlap of segmented peroxisomes and mitochondria to quantify spatial overlap at object level (Fig. 1a, Supp. Fig. 1b). The combined use of these metrics, signal overlap, correlated intensity patterns, and object-level spatial association, thereby provided a robust and nuanced assessment of the colocalization of mitochondria with peroxisomes and peroxisomal proteins.

To determine whether PEX14 was specifically enriched at mitochondria, we quantified the degree of colocalization between peroxisomal and mitochondrial markers. We found significantly higher colocalization of mitochondria with PEX14 than of mitochondria with PMP70 or ACAA1. Specifically, the MCC indicated a large fraction of total PEX14 signal (48%) overlapping with the mitochondrial signal, whereas only minor fractions of the PMP70 and ACAA1 signals (32% and 34%) colocalized with mitochondria (Fig. 1c). While a certain degree of colocalization is expected to occur randomly from spatial proximity of organelles, this effect equally affects all analyzed markers. Therefore, the substantially higher MCC observed for PEX14, corresponding to an increase of ∼15 percentage points compared to PMP70 and ACAA1, demonstrates a specific enrichment of PEX14 at mitochondria. This interpretation was reinforced by the significantly higher PCC observed for PEX14 relative to the other peroxisomal markers, indicating a stronger global correlation between PEX14 and mitochondrial fluorescence intensity distributions (Fig. 1d).

Colocalization can result from physiological MCS between peroxisomes and mitochondria, which are formed by a mature population of peroxisomes. This mature population is defined by the presence of peroxisomal membrane proteins and the efficient import of cytosolic proteins into the peroxisomal matrix^13,41^. Accordingly, these peroxisomes are expected to be positive for PEX14, PMP70, and ACAA1. There is, however, no evidence for a PEX14 enrichment specifically in peroxisomes engaged in mitochondria - peroxisome contact sites. To nonetheless rule out that an apparent enrichment in protein abundance, reflected by higher fluorescence intensities, could confound our colocalization analysis, we determined the SOC. This approach quantifies spatial overlap based on object area, allowing assessment of colocalization independently of fluorescence intensity. Using this method, we found that 12% of the segmented PMP70-signal overlapped with the mitochondrial area. In contrast, a significantly larger area of 18%, segmented by the PEX14-signal, was associated with mitochondria (Fig. 1e). This higher overlap demonstrates an additional mitochondrial association of PEX14 that cannot be explained by peroxisome - mitochondria contact sites.

As a control, COX IV, a protein of the inner mitochondrial membrane (IMM), expectedly localized along mitochondrial tubules and showed a high degree of colocalization with mitotracker using established metrics (Fig. 1c, d) and object-based colocalization (Fig. 1e).

Our quantitative analysis of more than 600 cells identified a previously underappreciated mitochondrial localization of endogenous PEX14 in human cells. This mitochondrial PEX14 fraction is sufficiently abundant to be reliably detected by high-content confocal fluorescence microscopy, supporting the conclusion that endogenous PEX14 localization at mitochondria is a reproducible and quantifiable feature under physiological conditions.

To test if this finding was specific for U2OS cells, we investigated the endogenous localization of PEX14 in three additional human cell lines originating from lung (A549), kidney (HEK293), and cervix (HeLa) (Fig. 2, Supp. Fig. 2a, b). The largely uniform and ubiquitous distribution of peroxisomes without specific hotspots within the cytoplasm (PEX14 and PMP70 puncta) in these cells was similar to the distribution we found in U2OS cells. We identified regions in all cell lines where PEX14 and PMP70 are located in close proximity to mitochondria, as well as areas where these signals colocalize with mitotracker (Fig. 2a).

**Fig. 2:**
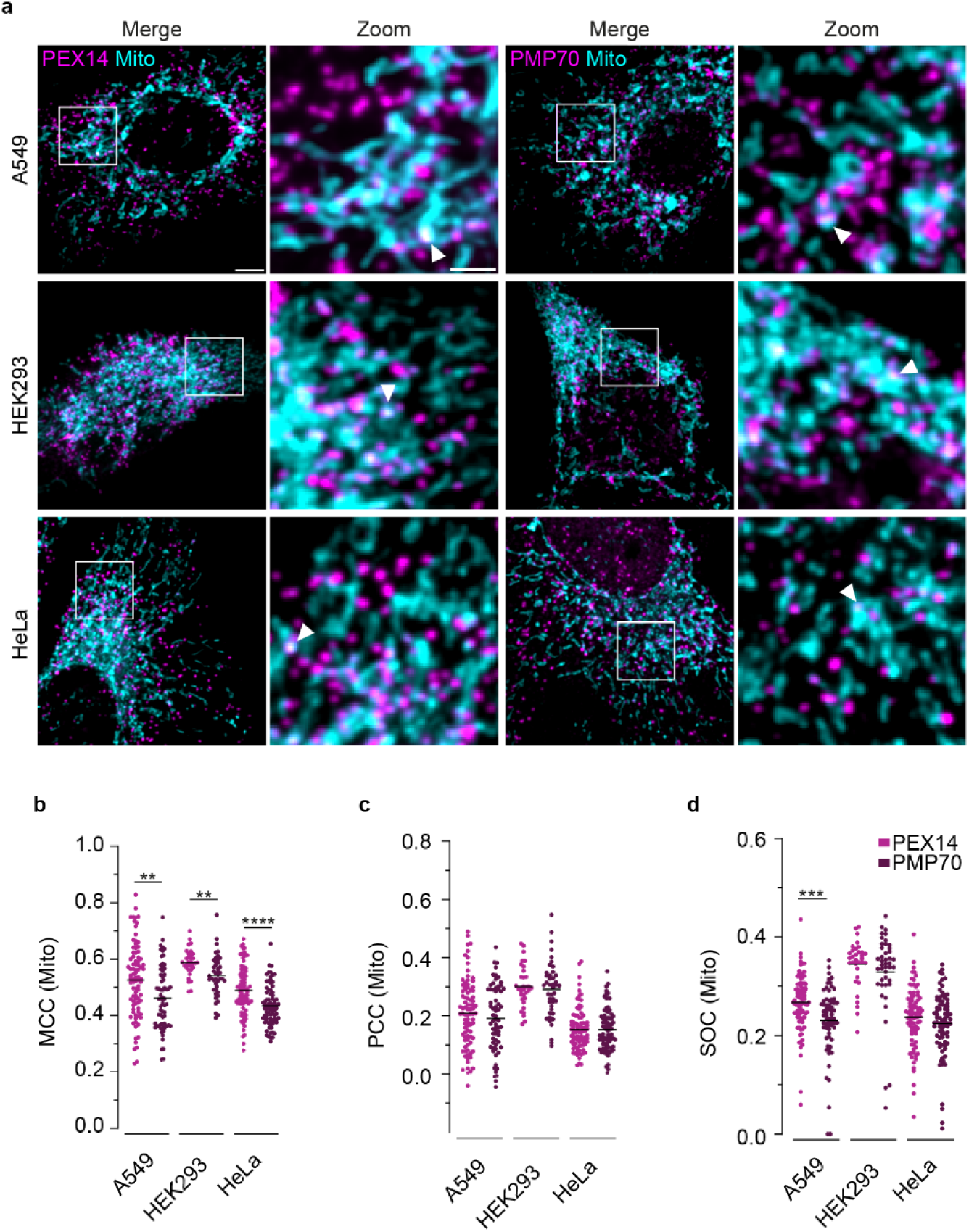
Endogenous PEX14 localizes at the mitochondrial membrane in various human cells. **a** Representative two-color confocal fluorescence microscopy images of different cell lines immunostained for endogenous PEX14 and PMP70 (magenta): A549 (N = 87 cells for PEX14, N = 66 cells for PMP70), HEK293 (N = 33 cells for PEX14, N = 43 cells for PMP70), and HeLa (N = 88 cells for PEX14, N = 82 cells for PMP70). Mitochondria were stained with mitotracker (cyan). Examples of colocalizing puncta are indicated by arrowheads. Scale bar: 5 µm. Zoom of the indicated box: scale bar: 2 µm. **(b-d)** Quantitative analysis of the colocalization of PEX14 and PMP70 with mitotracker. Cells are represented as individual points. The central line represents the median. **b** MCC with mitotracker. **c** PCC with mitotracker. **d** SOC with mitotracker. Statistical analysis was carried out using an unpaired t-test for PCC (A549, HEK293), MCC (A549, HEK293), and Mann-Whitney test for PCC (HeLa), MCC (HeLa), and SOC (A549, HEK293, HeLa): *P ≤ 0.05; **P ≤ 0.01; ***P ≤ 0.001; ****P ≤ 0.0001.

The extent of colocalization for all metrics varied across the different cell lines but was comparable to that observed in U2OS cells. Quantitative analysis of the different colocalization metrics indicates a stronger association of PEX14 with mitochondria than PMP70 for A549, HEK293, and HeLa cells (Fig. 2b-d). Significant enrichment of PEX14 at mitochondria was observed for the MCC in all cell lines (Fig. 2b). In addition, a significant difference in SOC was observed for A549 cells (Fig. 2d). Using a Wilcoxon signed-rank test applied to natural log-transformed colocalization ratios of PEX14 and PMP70 across all cell lines and metrics, we observed a significant ∼8% higher level of mitochondrial colocalization of PEX14 compared to PMP70 (Supp. Fig. 2c). Taken together, these quantitative analyses reveal the presence of endogenous PEX14 at mitochondria not only in U2OS WT cells, but in various human cell lines under physiological conditions.

### PEX14 localizes in the outer mitochondrial membrane

Since we showed the endogenous PEX14 localization at mitochondria by quantitative confocal fluorescence microscopy, it is not possible to distinguish between MCS and other PEX14-positive structures in proximity to mitochondria with this technique. To resolve all PEX14-containing structures, particularly those at or near mitochondria, we employed single-molecule localization microscopy. With dSTORM (direct stochastic optical reconstruction microscopy), it is possible to achieve a lateral resolution of ∼20 nm and to resolve individual PEX14 molecules. To gain insights into the distribution of PEX14 proteins within human cells, we analyzed localizations in more than 30 cells. We calculated minimal distances between individual localizations by k-nearest-neighbor (KNN) analysis and used a clustering algorithm to gain insights into the PEX14 distribution with respect to mitochondria (TOMM20, COX IV) (Fig. 3a).

**Fig. 3.**
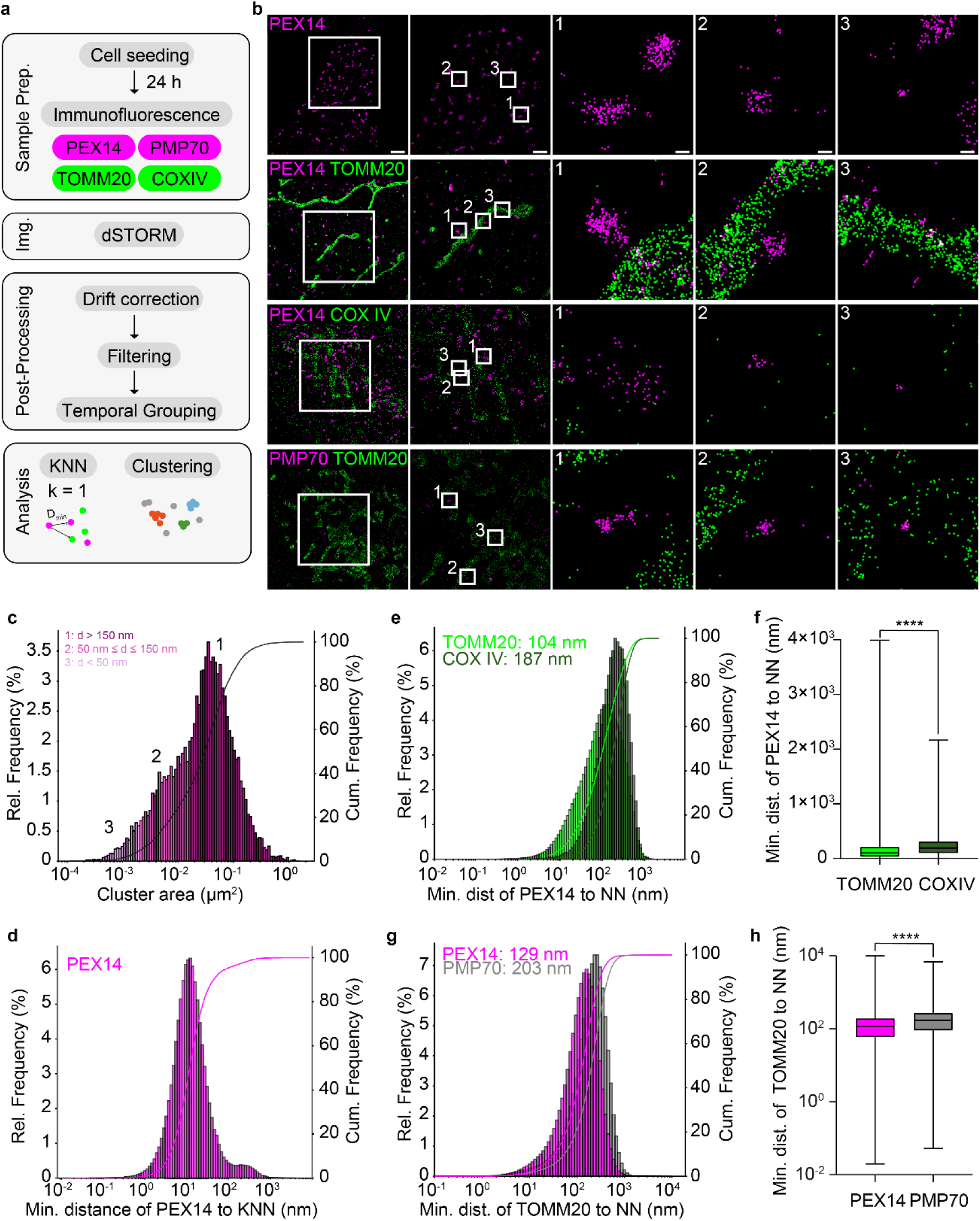
dSTORM reveals the localization of PEX14 in the outer mitochondrial membrane. **a** Workflow of sample preparation for dSTORM imaging, including subsequent post-processing consisting of drift correction, filtering, temporal grouping, and analysis of localizations using KNN and clustering. **b** Reconstructed two-color overview dSTORM images based on single-molecule localizations obtained after post-processing of representative cells. U2OS WT cells were immunostained for endogenous PEX14 or PMP70 (magenta) together with TOMM20 or COX IV (green) as mitochondrial markers. Scale bar: 2 µm. A magnified view of the indicated box is shown, scale bar: 1 µm. Three examples of the described PEX14 structures of different sizes, indicated by further boxes, were shown in Zoom: classes 1-3, scale bar: 100 nm. **(c-h)** KNN and clustering analysis of U2OS WT cells (N = 31 cells). At least three independent experiments were conducted. The histograms show the relative frequency (left y-axis) and the cumulative frequency (right y-axis), including a corresponding curve. **c** Distribution of PEX14 cluster size in µm^2^, divided into different populations based on cluster size determined by diameter: d >50 nm (1, light purple); 50 nm ≤ d ≤ 150 nm (2, purple), and d > 150 nm (3, dark purple). **d** Distribution of the minimum distances between PEX14 localizations (N> 1.7 × 10^6^ localizations) and their k = 1 nearest PEX14 neighbor in nm. **e** Distribution of the minimum distances between PEX14 localizations and their k = 1 nearest TOMM20 (light green) (N > 4.9 × 10^6^ localizations) or COX IV (dark green) (N > 0.77 × 10^5^ localizations) neighbor in nm. **f** Boxplot of the data shown in **(e)**. Statistical analysis was carried out using a two-sided Kolmogorov-Smirnov test with ****P ≤ 0.0001. **g** Distribution of the minimum distances between TOMM20 localizations and their k = 1 nearest PEX14 (magenta) or PMP70 (grey) (N > 1.0 × 10^6^ localizations) neighbor in nm. **h** Boxplot of the data shown in **(g)**. Statistical analysis was carried out using a two-sided Kolmogorov-Smirnov test with ****P ≤ 0.0001.

Profiting from the high resolution of dSTORM, we observed multiple forms of PEX14 signals based on their spatial distribution and size (Fig. 3b), while confocal microscopy only revealed a uniform distribution of punctate structures with similar sizes. To quantitatively assess these populations, we performed DBSCAN clustering of over 1.7 million PEX14 localizations and analyzed the resulting cluster areas, where clusters represent spatially dense groups of localizations (Fig. 3c). We identified the presence of different PEX14 populations by using a Gaussian mixture model (GMM) (Supp. Fig. 3). The resulting size distribution was broad, with a median of ∼0.028 µm^2^ and a range spanning ∼0.0001 to 1.6 µm^2^, indicating substantial diversity in the distribution of PEX14 molecules.

Based on the observed heterogeneity of PEX14 signal, we defined three classes of PEX14-positive structures: large, round or oval structures with diameters >150 nm and high localization density (class 1); intermediate-sized structures with diameters of 50 - 150 nm with fewer localizations but a similarly round appearance (class 2); and small accumulations of PEX14 within diameters <50 nm (class 3) (Fig. 3b). The size of the largest structures (class 1) is consistent with reported peroxisome dimensions^42^, suggesting that many of these represent mature peroxisomes. In contrast, structures within the 50 - 150 nm (class 2) range fall within the typical size reported for MDVs^43,44^ and ppVs^33^, indicating that these structures are likely vesicular intermediates. Structures with a diameter of 50 nm or smaller (class 3) represent smaller assemblies, including individual PEX14 oligomers, whose apparent size may appear enlarged due to antibody linkage error.

While the majority of PEX14 localizations (63%) were associated with large clusters corresponding to peroxisomes (class 1), a considerable fraction (32%) was assigned to vesicular structures (class 2). Additionally, a small subpopulation (5%) was found in clusters with diameters <50 nm (class 3), consistent with small structures, like oligomeric assemblies. Our analysis revealed that a large proportion of 37% (classes 2 and 3) of all PEX14 molecules are organized within structures that cannot be resolved by confocal microscopy, further highlighting the necessity of high-resolution microscopy for the study of PEX14.

While clustering helped determine the overall size and distribution of PEX14-positive structures, it does not provide information on local spatial relationships between individual localizations. Therefore, we next applied nearest-neighbor analysis. The majority (95%) of PEX14 molecules were located within 100 nm of the next PEX14 signal, with a median distance of 11.8 nm (Fig. 3d, Supp. Table 3). Given that PEX14 is part of the oligomeric peroxisomal matrix protein import complex, such short intermolecular distances can be expected^45,46^. These small distances highlight the necessity of super-resolution microscopy to accurately resolve PEX14 organization at nanoscale. However, as this analysis does not distinguish between peroxisomal and mitochondrial PEX14, similarly short distances between PEX14 molecules may also arise in mitochondrial or vesicular membranes.

Therefore, we investigated the mitochondrial localization of PEX14 by co-staining with mitochondrial markers. To assess whether PEX14 localizes to the outer or inner mitochondrial membrane, membranes that are separated by ∼6-12 nm^47^ and cannot be resolved by conventional confocal microscopy, we used TOMM20 as a marker for the outer mitochondrial membrane (OMM) and COX IV for the IMM (Fig. 3b).

Focusing on TOMM20, we observed that peroxisome-type large PEX14-positive structures (class 1) were not only dispersed throughout the cytoplasm but were also frequently observed in the proximity to the OMM, suggesting the presence of mitochondria - peroxisome contact sites. Vesicular-type intermediate-sized PEX14 clusters (class 2) also showed associations with the OMM, indicating that such interactions are not restricted to the larger-sized peroxisomes. Of note, a substantial fraction of small PEX14 clusters (class 3) was found interspersed within TOMM20 localizations, suggesting their integration into the OMM (Fig. 3b, TOMM20). Staining with COX IV showed less association of PEX14 with the IMM. However, PEX14 and COX IV structures with close contact to each other can also be found here, which could be explained primarily by MCS for the larger PEX14 structures (class 1) (Fig. 3b, COX IV).

To confirm the localization of PEX14 in one of the mitochondrial membranes, we calculated the distance of each PEX14 localization to the nearest TOMM20 or COX IV neighbor (k = 1) with nearest-neighbor analysis (Fig. 3e, f; Supp. Table 4-5). The significantly smaller distance of PEX14 to TOMM20 (median 104 nm) compared with the distance of PEX14 to COX IV (median 187 nm) indicates that PEX14 is located in the OMM, not the IMM.

Together, these results demonstrate that PEX14 is distributed across multiple structural populations, all of which can be associated with mitochondria. The smallest PEX14-positive structures are observed in the OMM but not in the IMM.

Since peroxisomes are known to physically associate with mitochondria, proximity to TOMM20 alone could reflect organelle-level contact rather than a protein-specific distribution. To distinguish between these possibilities, we co-stained PMP70 with TOMM20 (Fig. 3b) and compared the TOMM20 - PEX14 distance to that of TOMM20 and PMP70 (Fig. 3g, h; Supp. Tables 6-7). The significantly greater distance to TOMM20 observed for PMP70 (203 nm vs. 129 nm for PEX14) demonstrates that mitochondrial proximity is not a general feature of peroxisomal membrane proteins through MCS between peroxisomes and mitochondria but is selectively associated with PEX14. This is consistent with the enriched mitochondrial localization of PEX14 detected by confocal imaging (Fig. 1).

In conclusion, dSTORM confirmed mitochondrial localization of endogenous PEX14 and, in addition, demonstrated that PEX14 resides in close proximity to TOMM20 in the OMM. The localization of endogenous PEX14 in the mitochondrial membrane in WT human cells may suggest a role of this PEX14 subpopulation in the de novo biogenesis of peroxisomes from mitochondria.

To independently examine endogenous proteins at mitochondria, we performed unbiased spatially resolved proteomic analysis using only endogenous TOMM20 as a marker (Fig. 4a,b). This approach allowed us to analyze the proteomic composition of regions in the vicinity of the OMM independently of any peroxisomal marker. Mitochondrial regions of interest (ROIs) were identified using TOMM20 immunostaining prior to photoactivation of the proximity probes (Fig. 4b,c). Biotinylated proteins of the photolabeled ROIs (Supp. Fig. 4a) were subsequently isolated and analyzed by mass spectrometry.

**Fig. 4:**
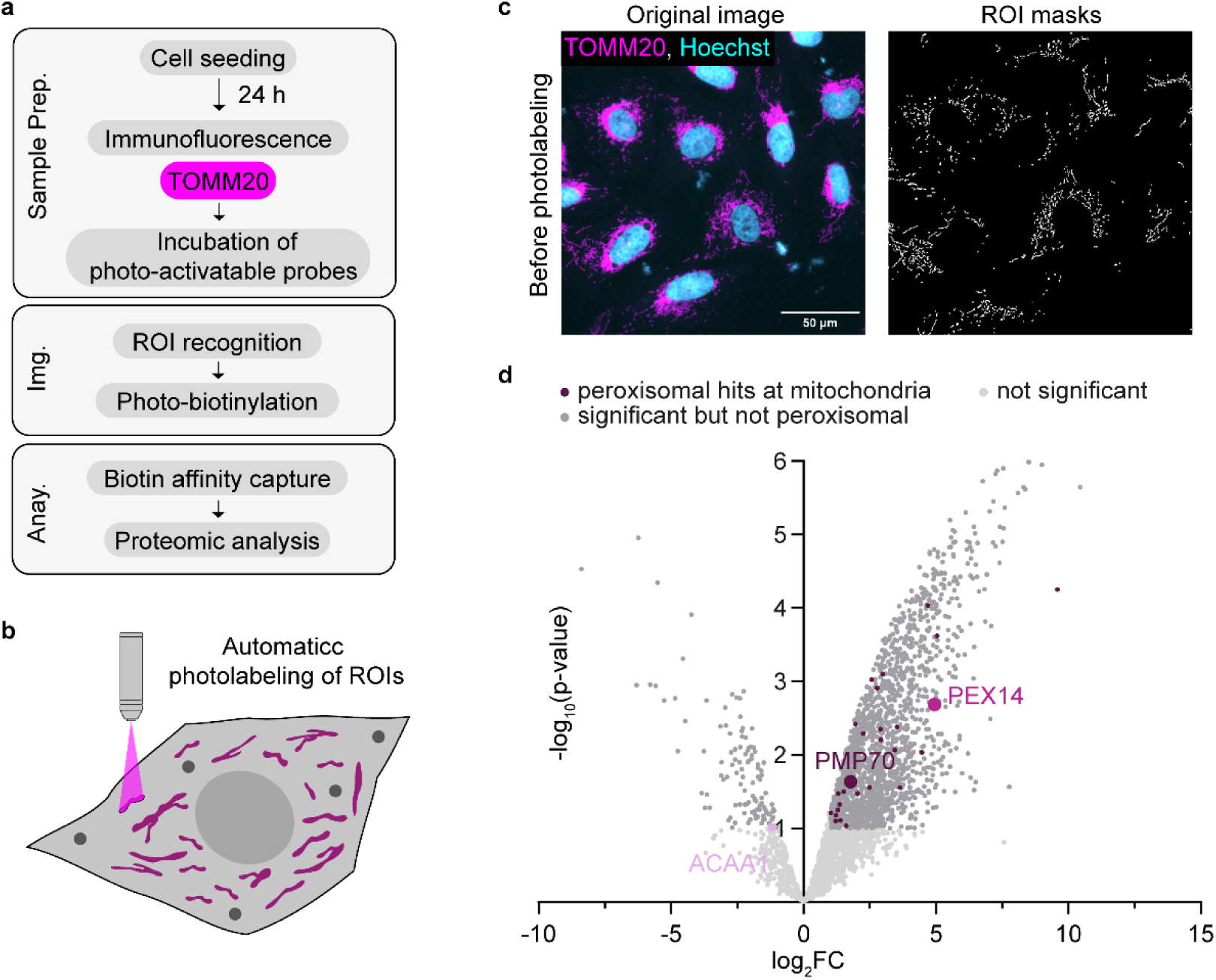
Spatial proteomics demonstrates mitochondrial enrichment of PEX14. **a** Workflow of sample preparation for spatial proteomics, including IF of endogenous TOMM20, incubation with photo-activatable probes, ROI recognition and photo-biotinylation, followed by biotin affinity capture and proteomic analysis. **b** Schematics of mitochondrial ROI recognition and photolabeling, not to scale. **c** U2OS WT cells before photolabeling immunostained for TOMM20 (magenta) and Hoechst (cyan) as a nuclear marker with ROI masks used for subsequent photolabeling. **d** Volcano Plot of photolabeled versus unlabeled U2OS WT cells immunostained for endogenous TOMM20. All significantly enriched proteins, exhibiting a known peroxisomal but non-mitochondrial localization, were shown in violet. PEX14 (FC = 30.9) is enriched at mitochondria to a much higher extent than PMP70 (FC = 3.5) or ACAA1 (FC = 0.45).

More than 2,000 proteins were identified in mitochondrial ROIs, including 41 peroxisomal proteins. Among these, PEX14 showed significant enrichment at mitochondria, whereas other peroxisomal proteins were also detected at the OMM but showed substantially lower enrichment, with PEX14 enrichment being ∼ 8-fold higher than that of PMP70 and more than 60-fold higher than that of ACAA1 (Fig. 4d). This enrichment of PEX14 could not be explained by a higher overall protein abundance compared with PMP70 or ACAA1, as PEX14 is not expressed as higher levels than PMP70 or ACAA1 (Supp. Fig. 4b). Thus, the preferential detection of PEX14 in proximity to TOMM20 is unlikely to result from differences in overall protein abundance but rather supports the specific association of endogenous PEX14 with mitochondria.

Together with our imaging data, this orthogonal approach demonstrates the endogenous PEX14 localization at mitochondria in human cells under physiological conditions.

### PEX14-specific mitochondrial localization

To investigate a potential role of PEX14 at mitochondria, we imaged living cells with various HaloTag (HT)^48^ constructs. This allowed us to observe the in vivo localization of PEX14 and, at the same time, avoid fixation artifacts. U2OS cells were transfected, and after 24 h, the expressed protein was labeled for 1 h using HT ligand. Mitochondria were co-stained in parallel using mitotracker (Fig. 5a).

**Fig. 5:**
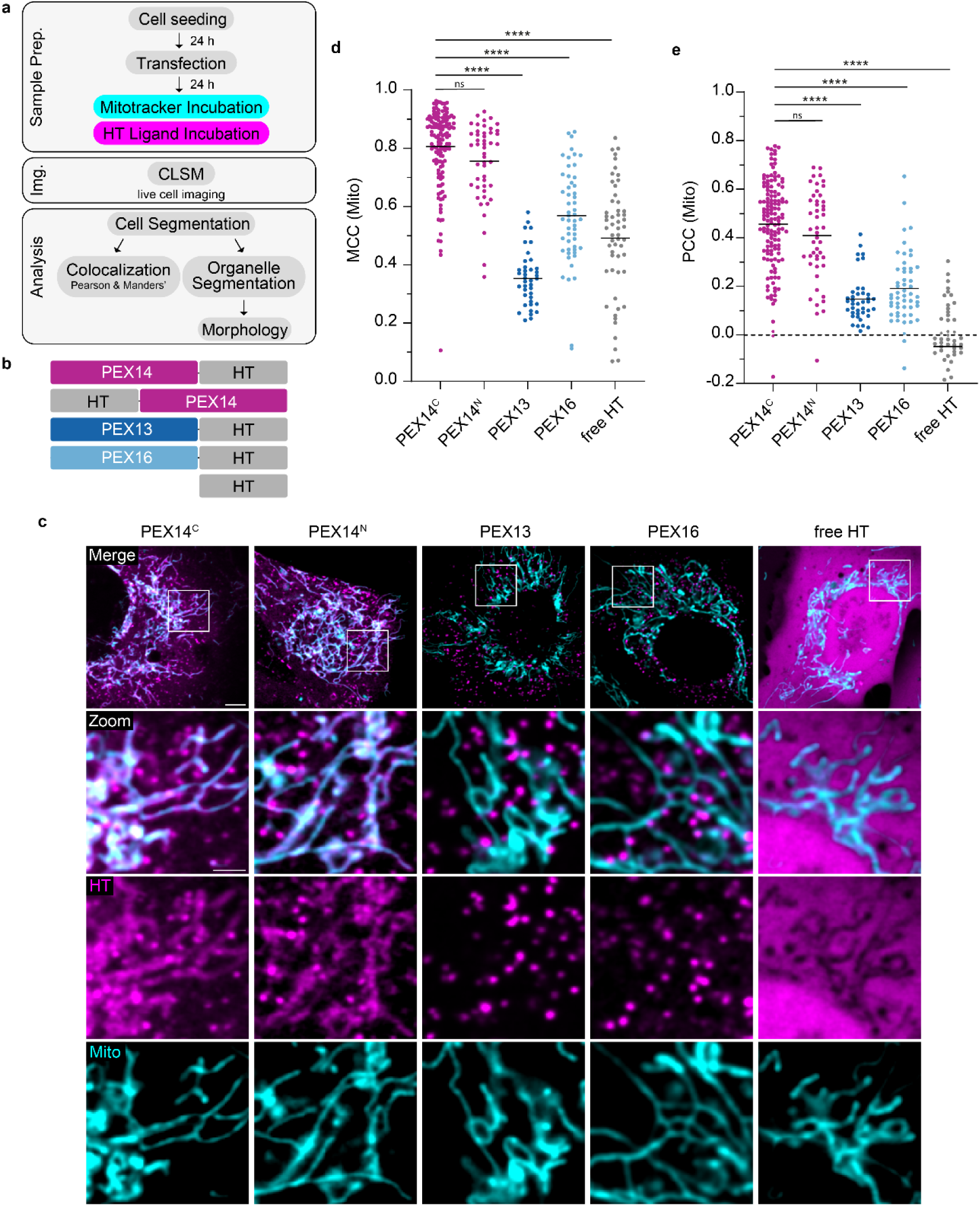
PEX14-specific mitochondrial localization upon overexpression. **a** Workflow of sample preparation for live cell imaging, including subsequent analysis of cells and organelles. U2OS WT cells were transfected 24 h before incubation with mitotracker and HT ligand. Cells were imaged at 37°C and 5% CO_2_. **b** Schematic of the transfected PEX constructs; not to scale. **c** Representative two-color live confocal fluorescence microscopy images of U2OS WT cells expressing the different HT constructs (magenta) co-stained with mitotracker (cyan). Scale bar: 5 µm. Zoom of the indicated box: scale bar: 2 µm. **(d-e)** Quantitative analysis of the colocalization of PEX14^C^ (N = 136 cells), PEX14^N^ (N = 48 cells), PEX13 (N = 40 cells), PEX16 (N = 53 cells), and free HT (N = 55 cells). At least three independent experiments were conducted. Cells are represented as individual points. The central line represents the median. Statistical analysis was carried out using the Kruskal-Wallis test with two-sided Dunn’s multiple comparison test: ns > 0.05; *P ≤ 0.05; **P ≤ 0.01; ***P ≤ 0.001; ****P < 0.0001. **d** MCC with mitotracker. **e** PCC with mitotracker.

In order to contextualize the localization of exogenous PEX14 expression, we compared it with that of two other peroxisomal membrane proteins involved in peroxisome biogenesis. PEX13 is present in the peroxisomal membrane as part of the import complex, together with PEX14^34,49,50^. We also used PEX16, as it plays an important role in PMPs topogenesis in peroxisome membrane formation^51–53^ and is a key player in de novo biogenesis as part of ppVs derived from the ER^39^. All peroxins were C-terminally tagged. Additionally, we N-terminally tagged PEX14 to assess a possible influence of the tag position on localization. As a control, we used the free HT (Fig. 5b).

We examined the intracellular distribution of the HT signal within the cell and observed dual localization for both C- and N-terminally tagged PEX14 variants. In addition to abundant PEX14-positive puncta, which we assume to be peroxisomes or vesicles based on their size and morphology, we also detected PEX14 at mitochondria, consistent with previously reported findings^24^ (Fig. 3c). In contrast, PEX13 and PEX16 showed an exclusively punctate signal with no mitochondrial localization.

As previously discussed, recombinant expression of peroxins can interfere with peroxisomal dynamics and lead to altered localization, a phenomenon often referred to as mistargeting. However, PEX13 and PEX16 retained their expected punctate, likely peroxisome-associated, localization, which does not colocalize with mitotracker, demonstrating that overexpression of peroxins does not generally lead to mistargeting to mitochondria. The selective mitochondrial localization of PEX14, therefore, cannot be attributed to a non-specific overexpression artifact but rather supports the specific association between PEX14 and mitochondria. Consistent with this interpretation, the free HT control displayed a diffuse cytoplasmic distribution (Fig. 4c), confirming that the tag itself does not drive organelle association.

This observation was supported by colocalization measurements with mitochondria (Fig. 5d,e). Both PEX14 constructs showed significantly higher MCC values of 0.84 and 0.81, respectively, than PEX13 (0.35), PEX16 (0.56), and the free HT (0.53) (Fig. 5d). These observations were confirmed when we determined the PCC (Fig. 5e). As localizations of the C- and N-terminally tagged PEX14 were indistinguishable, all subsequent experiments were conducted with the C-terminally tagged PEX14 construct. Colocalization values for PEX13 and PEX16 were in the same range as those for PMP70 and ACAA1 (Fig. 1c, d), supporting the notion that the respective localizations represent mature peroxisomes.

In conclusion, through imaging of living WT cells, we found that PEX14 is specifically localized to mitochondria. We did not observe a general mitochondrial localization of overexpressed peroxins but rather found this effect limited to PEX14.

In addition to affecting localization, expression of the various constructs can also influence other cellular processes and dynamics. High exogenous expression of proteins can induce cellular stress and lead to mitochondrial fragmentation^24^. Cells with high exogenous protein expression typically require lower laser power for imaging. By using a range of laser intensities, we demonstrated that higher PEX14 expression levels were associated with increased mitochondrial fragmentation (Supp. Fig. 5a). This fragmentation is characterized by an increase in mitochondrial roundness, as well as a reduction in the average mitochondrial area and length (Supp. Fig. 5b-d). To minimize potential effects of exogenous expression on mitochondrial dynamics, we used imaging conditions that ensured low to moderate expression levels of the different constructs.

### Mitochondrial PEX14 supports peroxisome biogenesis

To investigate if the subcellular localization of PEX14 affects peroxisome biogenesis, we used a series of PEX14 targeting variants. In addition to free HT as a control and the wild-type PEX14 sequence, we generated a primarily mitochondrial variant by fusion with the N-terminal mitochondrial targeting sequence (MTS) of TOMM20. Similarly, we used the membrane peroxisomal targeting sequence (mPTS) of PMP22 for a primarily peroxisomal localization of PEX14, even upon overexpression (Fig. 6a).

**Fig. 6:**
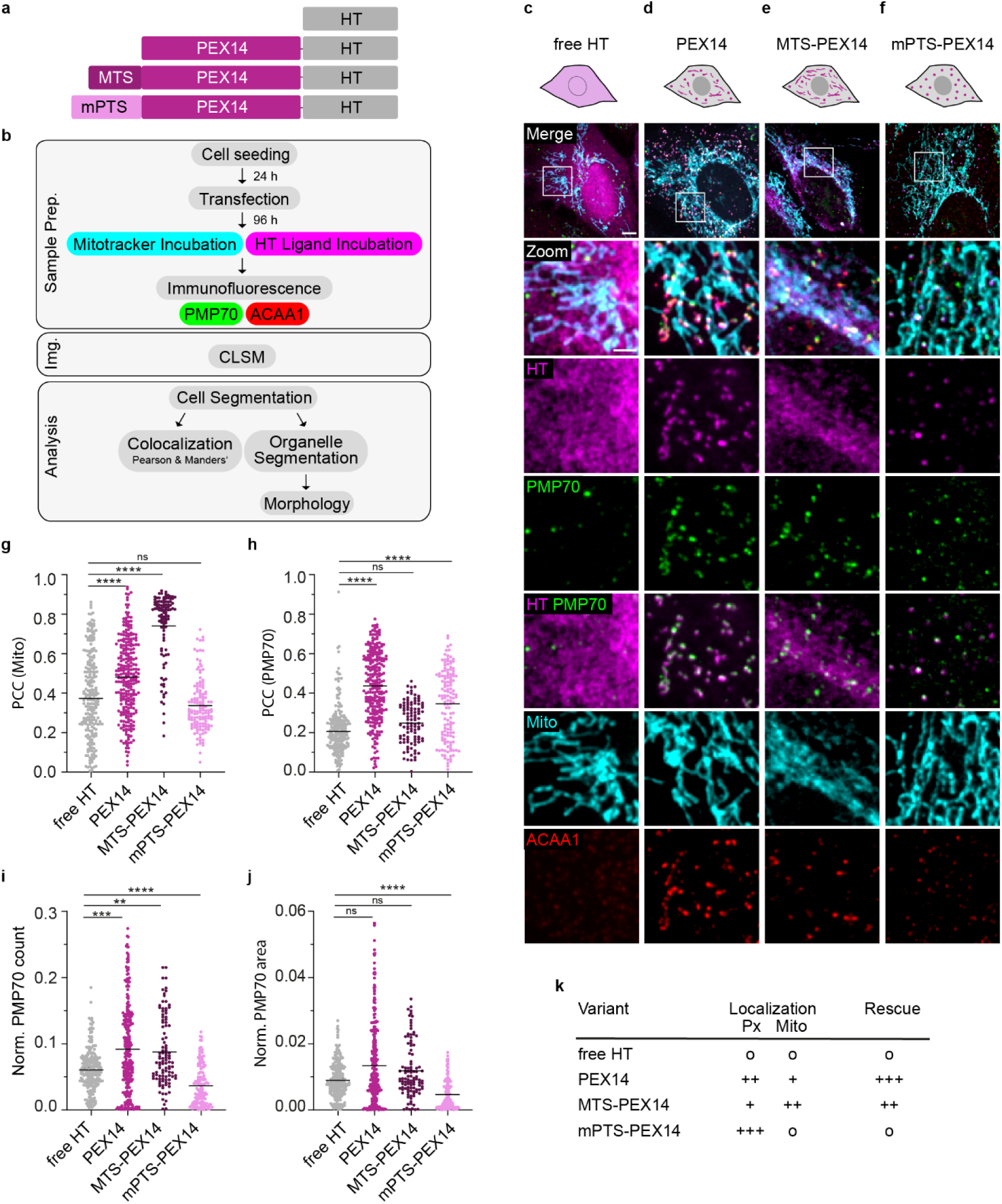
Mitochondrial PEX14 supports peroxisome biogenesis. **a** Schematic of the transfected PEX14 variants, not to scale. PEX14 is shown in magenta, MTS in dark purple, mPTS in light purple, and HT in grey. **b** Workflow of sample preparation for confocal imaging, including subsequent analysis of cells and organelles. U2OS PEX14 KO cells were transfected 96 h before incubation with mitotracker and HT ligand, followed by immunostaining of PMP70 and ACAA1. **(c-f)** Representative four-color confocal fluorescence microscopy images of U2OS PEX14 KO cells expressing the different PEX14 variants (magenta). Schematic subcellular localization is shown in the upper panel. Cells were co-stained with mitotracker (cyan), endogenous PMP70 (green), and endogenous ACAA1 (red). Scale bar: 5 µm. Zoom of the indicated box: scale bar: 2 µm. **c** free HT. **d** PEX14-HT. **e** MTS-PEX14-HT. **f** mPTS-PEX14-HT. **(g-h)** Quantitative analysis of cells expressing the different PEX14 variants (free HT N = 237 cells, PEX14 N = 263 cells, MTS-PEX14 N = 105 cells, mPTS-PEX14 N = 137 cells). At least three independent experiments were conducted. Cells are represented as individual points. The central line represents the median. Statistical analysis was carried out using the Kruskal-Wallis test with two-sided Dunn’s multiple comparison test: ns > 0.05; *P ≤ 0.05; **P ≤ 0.01; ***P ≤ 0.001; ****P ≤ 0.0001: **g** PCC with mitotracker as mitochondrial marker. **h** PCC with PMP70 as peroxisomal marker. **i** Normalized PMP70 count (number of PMP70 puncta per cell area in µm^2^). **j** Normalized PMP70 area (total PMP70 area per cell area). **k** Table of transfected PEX14 variants with their observed localization and rescue efficiency with o no, + weak, ++ intermediate, and +++ strong localization at peroxisomes/mitochondria and rescue effect, respectively.

We first confirmed the localization of each variant in U2OS WT cells (Supp. Fig. 6a-d). Free HT showed a cytosolic distribution (Supp. Fig. 6a), while wild-type PEX14 displayed its characteristic dual peroxisomal and mitochondrial localization (Supp. Fig. 6b), consistent with our findings in living cells (Fig. 5). The MTS variant localized predominantly to mitochondria (Supp. Fig. 6c), and the mPTS variant was found exclusively at peroxisomes, stained by PMP70 and ACAA1, with no detectable mitochondrial signal (Supp. Fig. 6d).

Notably, cells expressing MTS-PEX14 showed increased cytosolic ACAA1 signal and reduced ACAA1 import into PMP70-positive peroxisomes compared to free HT (Supp. Fig. 6c). We hypothesize that this is linked to the availability of cytosolic import receptors. Import of cargo carrying the peroxisomal targeting signal (PTS) into the peroxisomal lumen requires PEX14 and PEX13 together with the cytosolic receptor PEX5. For cytosolic proteins with a PTS-2, such as ACAA1, PEX7 is required additionally. After cargo binding of PEX5 in the cytosol, it binds to PEX14, allowing the cargo to be imported into the peroxisome through the PEX13 pore. Thus, binding of PEX5 to PEX14 is essential for the functional import of matrix proteins^10,35,49,50^.

Consistent with this, we found that overexpressed PEX14 recruited PEX5 not only to peroxisomes but also to mitochondria (Supp. Fig. 6e). In cells where PEX14 localizes predominantly to peroxisomes (Supp. Fig. 6e, cell 1), PEX5 colocalizes with PEX14 at peroxisomes and import proceeds normally. In cells with substantial mitochondrial PEX14 localization (Supp. Fig. 6e, cell 2), however, PEX5 is sequestered at the mitochondrial membrane, depleting the cytosolic pool of free PEX5 available for cargo binding. As a result, PTS-carrying proteins such as ACAA1 can no longer be efficiently captured in the cytosol, and their import into peroxisomes is impaired. Given that the formation of the functional importomer complex on mitochondria under pathological conditions has been shown^25^, the association of PEX5 with PEX14 at mitochondria was not surprising. We also noticed disruption of the import of PTS-1-carrying proteins when PEX5 was localized to mitochondria due to its binding to mitochondrial PEX14 (Supp. Fig. 6e, cell 2). Therefore, we assume that in cells expressing mitochondrial-localizing PEX14, endogenous PEX5 is bound to the mitochondrial membranes by the extensive amount of mitochondrial PEX14. Thus, PEX5 is not able to bind cytosolic cargo, like ACAA1, for peroxisomal matrix import.

To confirm that the targeting sequences do not affect general peroxisomal abundance, we compared PMP70 count and area in WT cells expressing each variant. While all PEX14 variants showed a minor reduction in peroxisome number and relative area compared to free HT, no differences were detected between the individual PEX14 variants (Supp Fig. 5f, g), suggesting that observed effects are attributable to PEX14 overexpression rather than the targeting sequences themselves. Together, these results indicate that the targeting constructs are functional. However, we cannot completely exclude the possibility that the functionality of PEX14 is influenced by the additional N-terminal targeting signal.

Additionally, we validated the native topology of PEX14, with its C-terminus exposed to the cytosol, using selective permeabilization of plasma- and intracellular membranes (Supp. Fig. 7). We analyzed the colocalization of the membrane-permeable HT ligand with the HT antibody (AB) for C- and N-terminally tagged PEX14 as well as for the MTS- and mPTS-PEX14 variants. Cells were either mildly permeabilized with digitonin, which permeabilizes the plasma membrane while leaving intracellular membranes largely intact, or fully permeabilized with Triton X-100, which disrupts cellular and intracellular membranes. As a control for peroxisomal membrane integrity under the digitonin conditions, we analyzed the luminal peroxisomal matrix protein ACAA1 (Supp. Fig. 7a). The colocalization between ACAA1 and the peroxisomal membrane protein PMP70 was reduced with digitonin compared with Triton X-100 treatment (Supp. Fig. 7c), demonstrating that the ACAA1 epitope was less accessible to the AB under mild permeabilization. This confirmed that the peroxisomal membrane was not permeabilized under the used digitonin conditions.

For N-terminally tagged PEX14, we observed less colocalization between the HT ligand and HT AB for digitonin compared with Triton X-100 treatment (Supp. Fig. 7b-c), indicating that the N-terminal HT epitope was not fully accessible from the cytosol. We therefore concluded that it was located within the peroxisome or mitochondrial intermembrane space. In contrast, no difference in HT ligand - HT AB colocalization between digitonin and Triton X-100 treatment was observed for C-terminally tagged PEX14, MTS-PEX14, or mPTS-PEX14 (Supp. Fig. 7b-c). For these constructs, the C-terminal HT epitope was accessible under digitonin conditions, revealing its cytosolic exposure. These results support the native C-out topology of all used PEX14 constructs.

Given their correct localization and functionality in WT cells, we wanted to assess peroxisome formation upon re-expression of PEX14 in the absence of endogenous PEX14. We generated two clonal CRISPR-Cas-mediated PEX14 knockout (KO) cell lines (Supp. Fig. 8a) and confirmed the absence of PEX14 (Supp. Fig. 8b-d). The PEX14 KO cells are characterized by a significant reduction in the number of peroxisomes (Supp. Fig. 8e, f). Due to the absence of PEX14, peroxisomal matrix proteins cannot be imported, resulting in the absence of mature, functional peroxisomes in PEX14 KO cells (Supp. Fig. 8e). All quantifications include data from both cell lines.

We transfected PEX14 KO cells with the PEX14-HT variants and ensured sufficient time (96 h) for peroxisome biogenesis to recover. Cells were then incubated with HT ligand and mitotracker for 1 h to visualize the PEX14 variants and mitochondria, followed by immunostaining of peroxisomes with antibodies against PMP70 and ACAA1. Using custom-written Python scripts for confocal analysis, we quantified colocalization and organelle morphology (Fig. 6b) based on segmentation of over 50,000 peroxisomes.

In PEX14 KO cells, each variant localized to its expected subcellular compartment, according to its localization in WT cells: free HT remained cytosolic (Fig. 6c), PEX14 showed dual peroxisomal and mitochondrial localization (Fig. 6d,g,h), MTS-PEX14 localized predominantly to mitochondria with markedly reduced peroxisomal signal (Fig. 6g,h) and mPTS-PEX14 was found exclusively at peroxisomes, with mitochondrial colocalization indistinguishable from free HT (Fig. 6f-h).

We next assessed the effect of the PEX14 variant re-expression on peroxisome formation in PEX14 KO cells by quantifying the normalized count and area of PMP70-positive structures, referred to as peroxisomes. Formation of new PMP70-positive peroxisomes was detectable upon re-expression of untargeted PEX14 and with more mitochondrial-localizing MTS-PEX14 (Fig. 6d,e). The normalized PMP70 count for these constructs increased from 0.058 for cells expressing free HT to 0.082 for PEX14 and 0.068 for MTS-PEX14 (Fig. 6i). These significant differences in normalized peroxisome count were supported by the visible trend of increased normalized PMP70 area (Fig. 6j). Interestingly, this rescue effect could not be demonstrated for mPTS-PEX14 (Fig. 6f), which was directly targeted to peroxisomes. In this case, no significant difference compared to the free HT was observed for either peroxisome count or area (Fig. 6i,j). These results indicate that mitochondrial PEX14 is necessary for the formation of new peroxisomes, since direct targeting of PEX14 to peroxisomes does not rescue the PEX14 knockout (Fig. 6k).

Strikingly, formed peroxisomes were consistently observed in close proximity to mitochondria, suggesting that peroxisomes may form at, or originate from, mitochondrial membranes. Possible differences in peroxisome formation observed upon re-expression of PEX14 in PEX14 KO cells are unlikely to result from variant-specific effects on steady-state peroxisome abundance, as no differences were observed in WT cells (Supp Fig. 5f, g), but instead reflect differences in de novo peroxisome biogenesis.

To further investigate peroxisome biogenesis following re-expression of the differently localized PEX14 variants, we additionally co-stained for the matrix protein ACAA1, since membrane integration of PMP70 alone does not represent the final step in peroxisome biogenesis and maturation. ACAA1 staining was therefore used to identify fully functional, import-competent peroxisomes. In PEX14 KO cells expressing free HT, no import-competent peroxisomes were present, as indicated by the absence of ACAA1-positive puncta colocalizing with PMP70 (Fig. 6c). In contrast, expression of PEX14 increased the number of ACAA1-positive puncta, which colocalized with PMP70 (Fig. 6d), demonstrating the restoration of import-competent, functional peroxisomes. Strikingly, import competence was similarly restored upon expression of MTS-PEX14 (Fig. 6e), demonstrating that mitochondrially targeted PEX14 supports peroxisome biogenesis.

We observed cells expressing PEX14 or MTS-PEX14 that failed to show a corresponding increase in ACAA1-positive puncta despite an increase in PMP70-positive peroxisomes. These cells exhibited higher HT signal intensities than cells displaying increases in both ACAA1 and PMP70 count. The mitochondrial localization of these PEX14 variants, together with our observation of PEX5 recruitment to mitochondria (Supp. Fig. 6e), suggests that this heterogeneity in import competence may result from limited availability of the import receptor. In line with this interpretation, higher HT intensities at mitochondria correlated with reduced ACAA1 import into peroxisomes, as reflected by a lower normalized ACAA1 count (Supp. Fig. 8g) and decreased colocalization of ACAA1 with PMP70 (Supp. Fig. 8h).

Strikingly, expression of the mPTS-PEX14 variant did not lead to any increase in ACAA1-positive puncta (Fig. 6f). This effect cannot be explained by sequestration of endogenous PEX5 at mitochondria, as this variant does not localize to mitochondria. Moreover, because no PMP70-positive peroxisomes were formed under these conditions, the absence of ACAA1-positive puncta further demonstrates that exclusively peroxisomal localization of PEX14 is insufficient to drive de novo peroxisome formation. Collectively, these results indicate that mitochondrial localization of PEX14 is required for efficient formation of peroxisomes.

We took into account that MTS-PEX14 predominantly localizes to mitochondria, but a fraction of the construct is also detected at peroxisomes. This is not unexpected, given that PEX14 itself is a peroxisomal membrane protein and contains information that can support targeting to and insertion into the peroxisomal membrane. Importantly, however, our conclusion that mitochondrial PEX14 supports peroxisome biogenesis does not rely solely on the localization of MTS-PEX14. Rather, it is supported by the functional comparison with mPTS-PEX14, which is restricted to peroxisomes. In PEX14-KO cells, expression of mPTS-PEX14 was not sufficient to restore the formation of new peroxisomes, whereas constructs localizing to mitochondria, such as MTS-PEX14, as well as PEX14 itself, supported peroxisome formation. Thus, although MTS-PEX14 is not exclusively mitochondrial, the inability of the peroxisome-restricted mPTS-PEX14 construct to restore peroxisome biogenesis indicates that mitochondrial targeting of PEX14 is required for efficient de novo peroxisome formation.

When combined with the endogenous localization of PEX14 to the outer mitochondrial membrane, these mechanistic analyses provide strong evidence that the mitochondrial subpopulation of PEX14 plays a functional role in the de novo biogenesis of peroxisomes.

## Discussion

In this study, we elucidate the role of PEX14 in peroxisome de novo biogenesis and reveal that its localization to mitochondria is critical for functional peroxisome formation. The model that mitochondria contribute directly to de novo peroxisome formation generated considerable interest^31^. However, a key open question has been whether endogenous peroxins physiologically localize to mitochondria or whether previously reported mitochondrial localization reflects mislocalization arising from peroxisome deficiency or protein overexpression^25^. In particular, it has remained unclear whether PEX14 naturally resides at mitochondria under physiological conditions. Therefore, we focused on the localization of endogenous PEX14 in human cells without perturbation of peroxisome biogenesis.

Using automated high-content confocal fluorescence microscopy imaging of human cells, we showed that PEX14 was enriched at mitochondria, in contrast to other peroxisomal markers such as PMP70 or ACAA1. This selective enrichment demonstrates that PEX14 is not exclusively associated with peroxisomes but naturally localizes to mitochondria under physiological conditions in human cells. Notably, mitochondrial enrichment of PEX14 was not restricted to U2OS cells but was consistently observed in three additional human cell lines and was independently shown by spatial proteomics of the OMM, supporting the existence of a conserved physiological mitochondrial pool of PEX14. Although conventional confocal fluorescence microscopy is sufficient to visualize peroxisomes and mitochondria as organelles, super-resolution microscopy is required to differentiate between peroxisomes, ppVs, and individual protein oligomers at different membranes. Whole-cell dSTORM imaging at 20 to 30 nm spatial resolution enabled us to resolve PEX14 molecules with nearest-neighbor distances of ∼10 nm and to identify distinct PEX14 clusters exhibiting pronounced size heterogeneity, with large, intermediate, and small structures. Notably, the smallest structures revealed the presence of PEX14 at mitochondrial membranes and, based on their spatial proximity to TOMM20 and COX IV, demonstrated the existence of a PEX14 subpopulation in the OMM. Intermediate-sized structures (diameter 50 - 150 nm) found in close proximity to mitochondria likely represent budding PEX14-positive MDVs. Consistent with our data, similarly sized PEX14-positive vesicles were recently reported in neurons^33^. Although the hybrid model of de novo peroxisome biogenesis postulates a localization of PEX14 at mitochondria, our data provide the first direct evidence that PEX14 endogenously localizes at mitochondrial membranes in human cells under physiological conditions, establishing mitochondria as a site of PEX14 function.

MDVs are a heterogeneous population that varies in membrane composition, cargo, and size depending on their function^32^. MDVs involved in peroxisomal interactions are frequently reported to be TOMM20-negative^44^. Intriguingly, a recent study identified TOMM20 as a component of ppVs^36^, whereas TOMM20-positive MDVs were more commonly associated with mitochondrial quality control^43,54^. These observations suggest that PEX14 associates with multiple, compositionally distinct MDV subtypes, underscoring the need to further define their heterogeneity and role in peroxisome biogenesis. Our data support a model in which ppVs arise from PEX14-positive mitochondrial membranes, indicating a PEX14-specific contribution to peroxisome biogenesis. However, because dSTORM relies on fixed-cell imaging, no conclusions can be drawn regarding the subsequent fate of these vesicles. In particular, how ER- and mitochondria-derived ppVs undergo fusion remains unclear. This question could be addressed in living cells using high-resolution imaging approaches such as single particle tracking (SPT).

To further define the role of PEX14 in de novo peroxisome biogenesis, we demonstrated a mitochondrial localization of PEX14 in living cells. Peroxin localization in mitochondria can occur upon loss of peroxisomes. For example, in a *pex19Δmsp1Δ* yeast strain, PEX14 and PEX13 were found highly enriched in mitochondria^25,55^. Given that both overexpression and loss of peroxisomes can potentially drive peroxins to mitochondria, a general co-enrichment of PEX13 and PEX14 at mitochondria might be expected. However, our analysis suggests that mitochondrial localization is limited to PEX14. Overexpression of PEX13 or PEX16 did not result in mitochondrial enrichment, also ruling out a general overexpression artifact with regard to PEX14. Together, these data suggest that the mitochondrial association of PEX14 is specific and reflects a functional relationship between PEX14 and mitochondria.

To directly test the functional importance of this mitochondrial localization for peroxisome biogenesis, we next employed a PEX14 KO background as a reductionist system in which PEX14 function could be dissected using defined variants with distinct subcellular targeting. Targeting sequences for the OMM and the peroxisomal membrane enabled us to demonstrate the importance of mitochondrial PEX14 in peroxisomal formation. Peroxisomes could only be restored upon re-expression of PEX14 variants that retained at least partial mitochondrial localization, as observed for both MTS-PEX14 and unmodified PEX14. Of note, neither construct localized exclusively to mitochondria, as both exhibited partial peroxisomal localization. In striking contrast, expression of PEX14 specifically targeted to peroxisomes failed to restore peroxisome formation.

A PEX14 KO, unlike a PEX3 or PEX19 KO, does not eliminate peroxisomes entirely. Accordingly, two scenarios could in principle account for the observed increase in peroxisome number and area: (i) restoration of pre-existing, non-functional peroxisomal membranes upon PEX14 integration, followed by their proliferation, or (ii) de novo formation of new peroxisomes. Our data argue against the first scenario. Expression of mPTS-PEX14, directly targeted to peroxisomal membranes, prevents peroxisome formation, demonstrating that a direct integration of PEX14 in non-functional peroxisomal membranes is not sufficient for restoration of peroxisome formation. In contrast, cells expressing PEX14 and MTS-PEX14, localized to mitochondria, recover peroxisomes efficiently, consistent with de novo biogenesis. In cells expressing PEX14 at low levels, where localization appears predominantly peroxisomal, we propose that a small fraction transiently localizes to mitochondria, likely below the detection threshold of confocal microscopy. We therefore favor a model in which new peroxisomes arise independently of pre-existing, dysfunctional membranes.

Using various microscopy techniques and quantitative analysis, we firmly established endogenous localization of PEX14 in the outer mitochondrial membrane. The specific mitochondrial localization upon PEX14 overexpression, and the necessity of this mitochondrial localization for peroxisome formation in PEX14 KO cells, reinforce the hypothesis that peroxisomes originate from PEX14-positive mitochondrial membranes.

PEX14 may be incorporated together with PEX3 into budding ppVs, which subsequently fuse with PEX16-positive vesicles derived from the ER. The resulting pre-peroxisomal compartment matures into a functional peroxisome through the sequential integration of additional peroxisomal membrane proteins, followed by the import of matrix proteins. In this framework, the E3 ubiquitin ligase MARCH5, which has been implicated in vesicle formation during de novo biogenesis and proposed to act upstream of PEX14, may regulate the mitochondrial budding step^36–38^. Another recent study identifies MARCH5 as a key player for mitochondria-driven quality control in peroxisome-deficient cells^56^. MARCH5 therefore represents an important target for future investigation. High-resolution live-cell imaging combined with endogenous protein tagging may help resolve the spatiotemporal coordination of ppV formation in living cells and further define the respective contributions of mitochondria and the ER to de novo peroxisome biogenesis.

## Material and Methods

### Molecular cloning

Plasmids generated in this study (Supp. Table 1) were constructed using standard molecular biology techniques, including PCR, restriction, and Gibson Assembly, and were verified by Sanger sequencing.

PEX14-HaloTag10 constructs were generated by amplification of HaloTag10, using Addgene plasmid #175525 and PEX14 (Gene ID: 5195) using mGFP-PEX14 via PCR, and either C- or N-terminal integration into the pEGFP-N1 vector backbone. For the generation of MTS-PEX14-HaloTag10, MTS (aa 1-33) of TOMM20^57^ was amplified via PCR using Addgene plasmid #175525. The mPTS (aa 1-133) of PMP22^58^ was amplified from HeLa cDNA via PCR for generation of mPTS-PEX14-HaloTag10. These targeting signals were cloned N-terminally to PEX14-HaloTag10 with a short 2x-Gly-Ser-linker.

The sequences for HaloTag9 and HaloTag11 for generation of PEX13-HaloTag9 and PEX16-HaloTag11 were amplified via PCR using Addgene plasmids #175525 and #175526. PEX13 (Gene ID: 5194) and PEX16 (Gene ID: 9409) were amplified from PEX13-mGFP and PEX16-mGFP. HaloTag sequences were fused C-terminally to PEX sequences with a 5x-Gly-Ser-linker and integrated into the pcDNA3.1(+) vector backbone.

Using PEX5-GFP, the sequence for PEX5 (Gene ID: 5830) was amplified. mNeonGreen sequence was amplified using Addgene plasmid #137804 and fused N-terminally to PEX5 with a 5x-Gly-Ser-linker integrated into pcDNA3.1(+) vector backbone.

For generation of pcDNA3.1(+)_mTagBFP2-PTS1, we used mTagBFP2, amplified from Addgene plasmid #55295, and cloned it N-terminally to PTS-1 containing the SKL (aa 691-700 of ACOX3 (Gene ID: 8310)).

### Cell culture and transfection

U2OS (American Type Culture Collection), HeLa (gift from Apl. Prof. Lübke, Bielefeld University), A549 (gift from Dr. Bednarz, Bielefeld University), and HEK293 (gift from Prof. Kaltschmidt, Bielefeld University) cells were cultured in high-glucose Dulbecco’s Modified Eagle Medium (DMEM) supplemented with 10% (v/v) fetal calf serum, 1% (w/v) L-glutamine, and 100 µg/mL penicillin/streptomycin (further referred to as supplemented DMEM) at 37°C and 5% CO_2_ and passaged using trypsin-EDTA. All cell culture media and supplements were purchased from Biowest.

Cells were transfected with plasmid DNA at a confluency of ∼75% using Lipofectamine 2000 transfection reagent (Invitrogen) for 24 hours according to the manufacturer’s instructions. For rescue experiments, cells were transfected 96 h before imaging. Total DNA amount was adapted according to cultivation area and cell number (100 ng for 8-well µ-slide, 1 µg for 12-well, 2 µg for 6-well). When co-transfecting multiple constructs, each construct represented an equal fraction of the total DNA amount.

For mitochondria staining, mitotracker variants were used: MitoTrackerRed CMXRos (Invitrogen) in experiments for fixed samples and MitoTrackerGreen FM (Invitrogen) or MitoSpy Orange CMTMRos (BioLegend) for live cell imaging. One hour before fixation or live imaging, respectively, mitotracker was added at a final concentration of 300 nM in supplemented DMEM and incubated at 37°C and 5% CO_2_.

When HaloTag constructs were transfected, Janelia Fluor JFX650 HaloTag Ligand (Promega) was added to the cells at a final concentration of 200 nM in supplemented DMEM for 1 h at 37°C and 5% CO_2_.

### Generation of PEX14 KO cell line

Clonal PEX14-KO cell lines were generated via CRISPR/Cas9-mediated gene editing. The CRISPR-RNAs (crRNA) targeting exon 1 of the human *PEX14* gene were designed as follows using CRISPOR^59^: crRNA1 5’-CCTCA GAAAG ATGGC GTCCT CGG-3’ and crRNA2 5’-AAAGA TGGCG TCCTC GGAGC AGG-3’ (Integrated DNA Technologies, IDT). By combining 100 µM crRNA with 100 µM trans-activating crRNA (tracrRNA), followed by heating to 95°C for 5 min, we formed functional guideRNA (gRNA) duplexes with a final concentration of 50 µM. Cells were nucleofected with gRNA duplex and Alt-R S.p.Hifi Cas9 Nuclease V3 (IDT) using SE cell line solution (Lonza) and U2OS cell-specific program CN-104 according to the manufacturer’s instructions, followed by single-cell sorting in 96-well plates 72 h after nucleofection. Clones were expanded, and genomic DNA (gDNA) was isolated using DirectPCR Lysis Reagent (Viagen Biotech). gRNA target sites were amplified by PCR with the following primers: 5’-GCGGT TGATT AGTCA GGCCT-3’ (OST3626) and 5’-ACAAG CTCAA GTGGG ACCAG -3’ (OST3627). PCR products were purified with Exo-CIP Rapid PCR Cleanup Kit (New England Biolabs, NEB) and Sanger sequenced (Supp. Fig. 8a). Two PEX14 KO clonal cell lines (PEX14 KO clone 1 and clone 2) were generated and verified with immunofluorescence (IF) and immunoblotting (IB) (Supp. Fig. 8b-d).

### Immunofluorescence

Cells were seeded in an 8-well µ-slide before staining and cultured as described previously. Subsequently, cells were washed three times with pre-warmed 1x phosphate-buffered saline (PBS) and fixed with 3.7% formaldehyde (FA) for 10 min at 37°C. All further steps were performed at room temperature (RT) unless otherwise noted. Permeabilization was typically done with 0.05% (v/v) Triton X-100 in PBS for 5 min. To investigate the topology of different PEX14-HT variants, cells were selectively permeabilized with 0.005% (v/v) digitonin in PBS for 5 min, rather than Triton X-100, to permeabilize the plasma membrane while preserving the integrity of the peroxisomal membrane. Non-specific binding sites were blocked with 5% (w/v) bovine serum albumin (BSA) in PBS for 30 min. Primary antibodies were diluted (Supp. Table 2) in 1% (w/v) BSA in PBS and incubated overnight at 4°C, followed by three washing steps with 1x PBS and incubation of secondary antibodies diluted 1:1,000 in 1% (w/v) BSA in PBS for 1 h at RT. Cells were washed three times with 1x PBS before further use. If nuclei were stained, 0.1 µg/mL DAPI was added during incubation of secondary antibodies. For confocal microscopy, cells were mounted in fluorescence mounting medium (Agilent).

### Confocal microscopy

Confocal fluorescence microscopy was done at RT using a laser-scanning confocal microscope (LSM900, Carl Zeiss Microscopy) with a 63x oil immersion objective (1.4 NA). (Immersol 518F/30°C, Carl Zeiss Microscopy). Images were acquired bidirectionally as 16-bit data with 4x averaging as z-sections, followed by maximum intensity projection using the ZEN 2 software (v 3.8, Carl Zeiss Microscopy) to generate a two-dimensional image.

### Live cell imaging

For imaging, the cell culture medium was changed to supplemented DMEM without phenol red. Cells were washed with this medium three times before imaging. Cells were imaged live at 37°C and 5% CO_2_ using a laser-scanning confocal microscope (LSM900, Carl Zeiss Microscopy) with a 63x oil immersion objective (1.4 NA) (Immersol 518F/37°C, Carl Zeiss Microscopy). Images were acquired as 16-bit data with 2x averaging at a single focal plane.

### dSTORM

For dSTORM imaging, cells were seeded in an 8-well µ-slide with a glass bottom (1.5 H) before fixation and cultivated as described before. Cells were washed once with pre-warmed 1x PBS and fixed with pre-warmed 3.7% FA for 10 min at 37°C. All further steps were performed at RT if not otherwise noted. Cells were washed with 1x PBS before incubation in reducing buffer (0.1% (w/v) NaBH_4_ in PBS) for 7 min. After washing three times with 1x PBS, cells were permeabilized with 0.1% (v/v) saponin in PBS for 5 min, washed once with 1x PBS, and blocked with 5% (w/v) BSA in PBS containing 0.1% (v/v) saponin for 30 min.

Primary antibodies were diluted (Supp. Table 2) in 1% (w/v) BSA in PBS containing 0.1% (v/v) saponin and incubated overnight at 4°C, followed by incubation with commercial Alexa Fluor 647 or Atto 488 conjugated secondary antibodies diluted 1:1,000 in 1% (w/v) BSA in PBS containing 0.1% (v/v) saponin for 1 h. After washing three times with 1x PBS, cells were postfixed with 3.7% FA for 10 min, followed by three final washing steps with 1x PBS.

Cells were imaged in Total Internal Reflection Fluorescence (TIRF) mode with an Oxford Nanoimager S (ONI) equipped with a 100x oil objective (1.45 NA) (Immoil-F30CC, Olympus) and sCMOS camera (Orca Flash4.0 v3, Hamamatsu) at the working temperature of the microscope (30 - 32°C). TetraSpeck microspheres (Thermo Fisher Scientific) were used for channel mapping calibration with the NimOS software (v1.19, ONI).

Immediately before imaging, 1x PBS was removed, and β-mercaptoethlylamine hydrochloride (MEA) buffer (200 µM MEA, 192 mg/mL glucose, with a glucose oxidase/catalase oxygen-scavenging system^60^, in 44 mM Tris-HCl, pH 7.0) was added to the cells. Two channels were imaged sequentially for 10,000 frames with 30 ms exposure (33.33 Hz), starting with 647 nm laser, before imaging with 488 nm laser.

### Spatial Proteomics

Spatial proteomics was performed using the Syncell Microscoop platform according to the manufacturer’s protocols for the Synlight-Rich kit (version 2.1) and SynPull kit (version 1.3), with adaptations described below. U2OS cells were seeded at 1.06 × 10⁵ cells per well in single-well chamber slides (Ibidi µ-Slide; 82106-90) precoated for 1 h at 37°C with Geltrex diluted in Opti-MEM (Gibco; 250 µL in 20 mL). Cells were cultured in DMEM (Gibco) with 10% FCS at 37°C and 5% CO_2_ for approximately 48 h and fixed at confluence with 4% paraformaldehyde containing 4% sucrose in PBS (pH 7.4) for 15 min at 37°C. Following three PBS washes, cells were blocked and permeabilized in PBS containing 10% goat serum and 0.3% Triton X-100 for 30 min. Mitochondria were stained with mouse anti-TOMM20 antibody for 1.5 h at RT, followed by Alexa Fluor 568-conjugated goat anti-mouse antibody for 1 h. Antibodies were diluted in blocking buffer, and samples were washed with PBS containing 0.3% Triton X-100 between incubations. Samples were subsequently treated with Block 1 and Block 2 reagents from the Synlight-Rich kit according to the manufacturer’s instructions. Nuclei were counterstained with Hoechst-JF646 (Janelia Farm; 1 µM) for 30 min at RT.

Microscopy-guided photobiotinylation was performed using the Microscoop platform (Syncell) and the Synlight-Rich photolabeling reagent, diluted 1:50 in PBS. Mitochondrial regions of interest were identified from TOMM20 fluorescence using Autoscoop software (version 1.0.1.11, Syncell) with perinuclear mitochondria excluded from the labeling mask. Imaging and photolabeling were performed on a Nikon Ti2-E microscope equipped with a Plan Apo Lambda D 40×/0.95 NA dry objective and a sCMOS camera (pco.edge 4.2) camera, using an 800 × 800-pixel acquisition format. TRITC and Cy5 images were acquired with 100 ms exposures at 10% and 5% LED intensity, respectively. Photolabeling was performed at 50% laser power with a labeling time of 100 µs and drift-free focusing. Three dishes were photolabeled, covering 12,040 fields of view per dish and approximately 0.9–1.1 × 10⁸ mask pixels per dish. Three corresponding unlabeled control dishes were processed in parallel, where the controls received photolabeling reagent but no photoactivation. After photolabeling, samples were washed and quenched in PBS containing 0.1% Triton X-100 using the kit reagent according to the manufacturer’s instructions. Labeling was verified by fluorescence imaging of reserved areas using the kit verification reagent.

Cells were harvested by scraping in SynPull collection buffer, and material from the three dishes was pooled separately for the photolabeled (PL) and unlabeled (UL) conditions. Protein extraction, enrichment, and digestion were performed using the SynPull kit (Syncell; protocol version 1.3). Samples were sonicated for 2 min, heated at 99°C for 1 h, treated with the kit’s subsequent lysis reagent for 30 min, and clarified by centrifugation at 16,000 × g for 20 min at 4°C. Protein concentrations were determined using the Pierce 660 nm protein assay with Ionic Detergent Compatibility Reagent (Thermo Fisher). An aliquot containing approximately 40 µg of protein from each pooled lysate was diluted in kit buffer and incubated with 5 µL of streptavidin magnetic bead suspension for 1 h at RT with mixing. Beads were washed using the SynPull wash buffers, including three final washes with 100 µL of Wash 4 buffer and brief sonication. Bound proteins were digested using the kit digestion reagent for 2 h, followed by overnight digestion for 16 h at 37°C. Digestion was stopped, and peptides were desalted using the kit reagents and desalting tips.

Peptides were analyzed by liquid chromatography–tandem mass spectrometry using an Orbitrap Astral Zoom mass spectrometer (Thermo Fisher) operated in data-independent acquisition mode. Each PL and UL sample, comprising material pooled from three dishes, was analyzed in three repeat LC–MS/MS injections, representing technical replicates. Raw data were processed using DIA-NN 2.3.2 against the UniProt Homo sapiens reference proteome (UP000005640; taxonomy ID: 9606), with contaminant sequences included. Searches used Trypsin/P specificity, up to two missed cleavages, fixed cysteine carbamidomethylation, variable methionine oxidation and protein N-terminal acetylation, and a 1% FDR threshold. Protein intensities were processed in R using a custom script, where contaminants were removed, and proteins detected in at least two of three technical injections in either condition were retained. Intensities were log₂-transformed, median-normalized, and imputed using MinProb (q = 0.01, tune.sigma = 1). PL and UL samples were compared using limma. Volcano plots displayed log₂(PL/UL) fold changes against −log₁₀(unadjusted P values), using thresholds of |log₂ fold change| > 1 and P < 0.1; positive fold changes indicated enrichment in the photolabeled fraction.

Relative protein abundance of PEX14, PMP70 and ACAA1 in U2OS WT cells was assessed with the PaxPb: Protein Abundance Database^61^ using all available U2OS WT proteomic datasets^62–64^.

### Immunoblotting

Cells were seeded in 6-well plates and cultured for 24 h as described previously before harvesting. After washing two times with ice-cold 1x PBS, cells were scraped into PBS. Via centrifugation at 14,000 rpm and 4°C, cells were pelleted and subsequently lysed in RIPA buffer (25 mM Tris-HCl, pH 8.0, 150 mM NaCl, 1% (v/v) NP-40, 1% (w/v) sodium deoxycholate, 0.1% (v/v) SDS) with 1x Pierce protease inhibitor (Thermo Fisher Scientific) by incubation for 30 min on ice. DNA was fragmented by multiple ultrasonic pulses in an ice-cold water bath for 10 min, and lysates were clarified by centrifugation for 15 min at 14,000 rpm at 4°C. Total protein concentrations were measured with a bicinchoninic acid (BCA) assay (Thermo Fisher Scientific).

Equal protein amounts of the samples were separated by SDS-PAGE and transferred to a 0.2 µm nitrocellulose membrane (BioRad). Membranes were blocked with 5% (w/v) milk in 1x TBS-T (Tris-buffered saline with 0.1% (v/v) Tween20) for 30 min at RT. Primary antibodies were diluted (Supp. Table 2) in 5% (w/v) milk in 1x TBS-T and incubated overnight at 4°C, followed by incubation of horseradish peroxidase (HRP)-coupled secondary antibodies diluted 1:10,000 in 5% (w/v) milk in 1x TBS-T for 1 h at RT. Blots were imaged with Clarity Western ECL Substrate (Bio-Rad) at ChemoStar Chemiluminescence Imager (Intas Science Imager Instruments). Band intensity values were measured using ImageJ (v 1.54p)^65^.

### Image analysis and visualization

Confocal images were processed and analyzed with a custom-written script (Confocal Microscopy Analysis) using Python (v 3.12). For segmentation of cells and organelles, images were pre-processed using normalization, Contrast Limited Adaptive Histogram Equalization (CLAHE), and Gaussian smoothing. Cells with no detected intensities were excluded from the analysis. Colocalization quantifications (MCC and PCC) were calculated using the raw data in ROIs defined as cells, segmented with Cellpose3 (cell diameter 35 to 45 µm, 10 retries)^66^. MCC values represent the fraction of the protein of interest (POI) signal colocalizing with mitochondria. Mitochondria and peroxisomes were segmented with Allen Cell and Structure Segmenter^67^. SOC was calculated by dividing the overlapping area of the segmented POI and mitochondrial signals by the total segmented POI area (Supp. Fig. 1b).

For experiments with transient transfection of constructs, a relative intensity threshold of 67% was set to only include transfected cells. To examine the effect of expression level, we calculated a relative expression by dividing the measured mean HT intensity within the cell by the product of the used detector gain and laser intensity.

For visual presentation of representative images, a Gaussian filter of 1 pixel was used, and brightness adjustments have been made with ImageJ (v 1.54p)^65^.

dSTORM data were post-processed and analyzed with a custom-written script (SMLM Analysis) using Python (v 3.12) previously described^68^. Identification of blinking events was done with the localization algorithm of NimOS (v 1.19, based on ^69^) using a point spread function (PSF) to localize a point-like object. Due to extensive blinking in the first 200 frames of each acquired channel, these frames were excluded from the analysis.

We drift corrected each channel individually, using adaptive intersection maximization (AIM)^70^ (segments of 100 frames, maximal distance of 20 nm, ROI radius of 60 nm), followed by alignment of the second acquired channel to the previous drift of the first acquired channel. To exclude artificial drift due to low blinking density often in the later frames, we defined a maximal drift threshold of 15 nm between two segments. If a calculated drift exceeded this value, the mean drift of the previous 10 segments was used for correction.

Drift corrected localizations were filtered using parameters as follows: Sigma of PSF 50 - 250 nm, localization precision < 20 nm, and p-value < 0.1 used in a chi-squared test of the log-likelihood of the fitted PSF. Only the localizations inside the top 95% percentile of photon count values were further processed. Localizations that occur within a maximal distance of 50 nm within a close timeframe (maximal frame gap of two) are grouped into a single event. An additional parameter for temporal grouping was the duration time of these localizations, between 1 and 50 frames. After post-processing and filtering, the median localization precision of the retained localizations was ∼10 nm.

Nearest neighbor analysis was conducted using the Euclidean metric with k = 1. Clustering was done using the Density-Based Spatial Clustering of Applications with Noise (DBSCAN) algorithm^70^ (ε = 100 nm, minPts = 10). To draw conclusions about the diameter of peroxisomes and vesicles, we assumed a perfect circle for the cluster shape, knowing that cluster shapes can vary.

### Statistical analysis

Statistical analysis was done with GraphPad Prism (v 9.1.1). Data was tested for normal distribution using the Shapiro-Wilk test. An unpaired two-sided t-test was used to compare two samples when the data were normally distributed with equal standard deviations (SDs). For data that was not normally distributed, a two-sided Mann-Whitney test was performed. The two-sided Kolmogorov-Smirnov test was used for non-parametric data. Comparing more than two datasets, we used Kruskal-Wallis test with two-sided Dunn’s multiple comparison test for non-normally distributed data. The statistical tests used for each dataset are indicated in the figure legend.

For global comparison across cell lines and colocalization metrics, a two-sided Wilcoxon signed-rank test against a theoretical median of 0 was used. The test was applied to natural log-transformed ratios of the mean colocalization values (PEX14/PMP70) to allow comparison across metrics with different scales and cell lines.

Distribution analysis was performed using Gaussian mixture modeling (GMM) implemented in Python (scikit-learn^71^). Models containing one to five Gaussian components were fitted using the expectation - maximization algorithm with full covariance matrices (random state = 42). Model selection was based on the Bayesian Information Criterion (BIC).

## Data Availability

Source data for all figures will be provided with this paper. Further data supporting the findings of this study are available from the lead contact upon request. Microscopy data reported in this paper will be shared by the lead contact upon request.

## Code Availability

SMLM Analysis was performed as previously described^68^. All original data analysis code is available in the provided Zenodo repositories: For Confocal Microscopy Analysis, see https://doi.org/10.5281/zenodo.20266296, and for SMLM Analysis, see https://doi.org/10.5281/zenodo.20268285. Additional information required to reanalyze the data reported in this paper is available from the lead contact upon reasonable request.

## Acknowledgments

We thank the Marlies und Herbert Repkow Stiftung for supporting the acquisition of an ONI Nanoimager and the DFG (TH 1538/3-1) and the Anschubfonds Medizinische Forschung (AMF) of Bielefeld University (projects AQP4x, RVtherapy, ERDKAR, and HC3P) for funding. We acknowledge the Core Facility Microscopy & Imaging for access to microscopes and a cell sorter. This work was supported by de.NBI Cloud within the German Network for Bioinformatics Infrastructure (de.NBI) and ELIXIR-DE (Forschungszentrum Jülich and W-de.NBI-001, W-de.NBI-004, W-de.NBI-008, W-de.NBI-010, W-de.NBI-013, W-de.NBI-014, W-de.NBI-016, W-de.NBI-022), which provided computing resources to conduct the analysis. We gratefully acknowledge Jana Heinrich for molecular biology work and Femke van Dijk for cloning and help with live-cell imaging. Furthermore, we thank Christina Heidar and Mona Pauli for help with selective permeabilization experiments. We thank Marcel Müller for insightful discussions on dSTORM post-processing and analysis workflow. We thank Matthias Plessner and the members of the Thoms lab for fruitful discussions. We thank Daniela Smoljanow and Simon Henning for comments on the manuscript.

## Author contributions

S.T. conceived and supervised the project and provided the funding. B.N. and A.S.H. designed and carried out all confocal, dSTORM, and functional experiments. B.N., A.S.H., and S.T. validated the data. Under the guidance of B.N. and A.S.H., S.K. developed the Python scripts for high-content and dSTORM image analysis. B.N. and A.S.H. analyzed and visualized the data. T.S., L.W. and M.L. performed the spatial proteomics experiments and analyzed the resulting data. B.N. wrote the initial draft of the manuscript. B.N. and S.T. edited and revised the manuscript with contributions from A.S.H., S.K., T.S. and M.L.

## Additional information

### Correspondence

Correspondence and requests for materials should be addressed to Sven Thoms.

